# Diols and sugar substitutes in attractive toxic sugar baits targeting *Aedes aegypti* and *Aedes albopictus* (Diptera: Culicidae) mosquitoes

**DOI:** 10.1101/2023.02.09.527878

**Authors:** Heidi Pullmann-Lindsley, Ava Bartlett-Miller, R Jason Pitts

## Abstract

Around the world, mosquitoes continue to transmit disease-causing pathogens while also developing resistance to insecticides. We previously discovered that a generally regarded as safe (GRAS) compound, 1,2-propanediol, reduces adult mosquito survivorship when ingested. In this study, we assess and compare five more chemically related compounds for mosquito lethality and eight GRAS sugar substitutes to determine toxic effects. We conducted a series of feeding assays to determine if ingesting the compounds influenced mosquito mean survivorship in locally collected lab-reared populations of *Aedes aegypti* (Linnaeus, 1762) and *Aedes albopictus* (Skuse, 1894) mosquitoes. Our results indicate that 1,2-propanediol, 1,3-propanediol, 1,5-pentanediol, 1,6-hexanediol, 2-methyl-1,3-propanediol, DL-dithiothreitol, acesulfame potassium, allulose, erythritol, sodium saccharin, stevia, and sucralose significantly reduced the mean survivorship of one or both species. Short-term trials with the most toxic compounds revealed that they could substantially affect survivorship after 24 hours. We also found that many of the compounds yielded different responses in the two species and that male mosquitoes expired to a greater extent than female mosquitoes. These findings indicate that several of the compounds may be highly effective for local mosquito vector population and disease control through functioning as components in attractive toxic sugar bait systems (ATSBs)

## Introduction

The frequency of insecticide resistance in mosquito disease vectors rises unrelentingly (Zaim and Guillet. 2002). Around the world, mosquito species have developed a variety of mechanisms to evade commonly used insecticides such as pyrethroids and organophosphates (Moyes *et al*. 2017, Fernando *et al*. 2020, Brogdon *et al*. 1992). Additionally, the concern for off-target effects on beneficial insects, including crucial pollinators such as *Apis mellifera*, grows (Goulson *et al*. 2015, Reid *et al*. 2020, Kruegar *et al*. 2021). These ongoing challenges encourage the development of novel control mechanisms.

One promising trend is integrating attractive toxic sugar baits (ATSB) into existing vector population control strategies (Fiorenzano *et al*. 2017). ATSBs typically combine a lure element, a sugar source, and a toxic component, attracting and eliminating adult mosquitoes (Frylewicz *et al*. 2021). Some commonly utilized features of ATSB construction include sucrose, fresh fruits or fruit juices, boric acid, and ivermectin (Frylewicz *et al*. 2021, Kumar *et al*. 2022, Scott-Fiorenzano *et al*. 2017, Tenywa *et al*. 2017). The usage of commonly available materials and ingredients, plus their low construction cost, make ATSBs an accessible option for local vector control in rural or developing areas (Stewart *et al*. 2013, Müller *et al*. 2010, Maia *et al*. 2018). Additionally, ATSBs include design elements that mechanically select against non-target species, such as a fine mesh or narrow opening, thus preventing larger insects from accessing the toxin or areas of application (Diarra *et al*. 2021, Qualls *et al*. 2015, Revay *et al*. 2015).

Numerous studies have demonstrated the efficacy of ATSBs in reducing populations of vector mosquitoes, including *Anopheles gambiae* (Giles 1902), *Culex quinquefasciatus* (Say 1823), *Ae. aegypti*, and *Ae. albopictus* (Traore *et al*. 2020, Qualls *et al*. 2015, Davis *et al*. 2021, Gu *et al*. 2020). In some treatments, the relative abundance of adult mosquitoes declined by as much as 90% (Müller *et al*. 2010). While these successes are encouraging, potential improvements can be envisioned. Notably, the components most often used do not explicitly target mosquitoes and could harm beneficial species (da Silva Cruz *et al*. 2010, Ferreira *et al*. 2013, Büyükgüzel *et al*. 2013). Therefore, the toxin is one area to investigate further in vector mosquito ATSB design. We previously validated 1,2-propanediol as a toxin against *Ae. aegypti, Ae. albopictus*, and *Cu. pipiens* under controlled laboratory conditions (Pullmann Lindsley *et al*. 2022). 1,2-propanediol is a generally-recognized-as-safe (GRAS) compound, meaning that it has been approved by the FDA for use in food and commercial products (Steele *et al*. 2016) and could be incorporated into ATSBs. Our results with 1,2-propanediol, in conjunction with other studies on the toxicity of diols to arthropods, led us to investigate other compounds for mosquito toxicity (Wang *et al*. 2021, Wegener *et al*. 2012).

In addition to investigating diols with chemical structures similar to that of 1,2-propanediol, we also evaluated a collection of compounds that share the GRAS status, specifically sugar substitutes. Each of the diols tested in this new study has commercial uses, such as in paints, personal care, hygiene products, cosmetics, and pharmaceuticals (Werle *et al*. 2008, Dionisio *et al*. 2018, Ren *et al*. 2018). Recent work demonstrates that certain sugar substitutes may be toxic to arthropods (Caponera *et al*. 2020, Barrett *et al*. 2020, Osabutey *et al*. 2021, Lee *et al*. 2021, Baudier *et al*. 2014, Rochlin *et al*. 2022); however, few artificial sweeteners have been studied for their toxicity to mosquitoes. Moreover, testing the non-nutritive sugar substitutes in combination with sucrose is critical to distinguishing between toxicity as a cause of death as opposed to starvation (Foster 1995, Barredo and DeGennaro 2020, Sharma *et al*. 2020).

To build upon our previous study (Lindsley *et al*. 2022) and add to our understanding of alternative mosquito toxins for ATSB design, we tested a group of fourteen compounds, comprised of diols and sugar substitutes, in a series of feeding assays for toxicity to two disease vector species, *Ae. aegypti* and *Ae. albopictus*. Based on our observations, we concluded that several compounds reduced survivorship in one or both species. These declines in vitality indicate that some of the compounds may be highly effective for local population control and could be effective against multiple mosquito species. We also observed differences in the toxicity of the compounds to *Ae. aegypti* and *Ae. albopictus*, as well as between males and females. Future studies will explore the effects of these compounds on non-target species, examine their efficacies in ATSB field trials, and assess other GRAS compounds, such as potassium sorbate and sodium benzoate, which have been used to control microbial growth (Neetoo and Mahomoodally 2014).

## Methods and Materials

### Mosquito Rearing

Larvae of *Aedes aegypti* and *Aedes albopictus* “McGregor” strains (Lindsley *et al*. 2022) were reared in a laboratory growth chamber under standard conditions: 27 °C, 70% RH, 12:12 LD in distilled water and were provided ground koi fish food mixed with baker’s yeast *ad libitum*. Pupae were collected and placed into cages (BugDorm-1; MegaView Sci. Co. Ltd., Taiwan) in clean distilled water prior to adult eclosion. Adults were allowed to mate for at least four days when females were separated and offered fresh defibrinated sheep blood (Hemostat Laboratories, Dixon, CA, USA), heated to 37°C using an artificial feeding system (Hemotek Ltd., Blackburn, UK).

### Ad libitum Trials

At the pupal stage, mosquitoes were divided into cages of approximately 50 individuals and provided 5% sucrose solution (w/v) in a 25 ml glass bottle with a cotton wick. Approximately seventy-two hours post-eclosion, the sucrose solution and pupal water cup were removed from the cages, and the adult mosquitoes were starved for twenty-four hours. The exact numbers of mosquitoes in each trial are provided in supplemental data. After starvation a 5% sucrose (w/v) plus 5% toxin (w/v) solution was provided in a 25 ml glass bottle with a cotton wick. One milligram of powdered dye was added to each sugar solution to monitor feeding status (Acid Blue 9, TCI America, CAS # 3844-45-9). Information about individual toxins is available in Supplementary Materials (Supp.Table S1). For control cages, mosquitoes received a 5% sucrose solution, dyed blue, in the same manner. Cages were visually inspected every 24 h for 7 days. Each day, expired mosquitoes were removed, sexed, checked for evidence of dye in the abdomen, and counted. At the end of the 7-day trial period, remaining mosquitoes were sexed and counted. Three biological replicates for each trial were conducted.

### 24-Hour Trials

Trials were initiated as described above. Mortality was assessed after 24 h, and the toxin was removed and replaced with a 5% sucrose solution, colored blue. Mortality was assessed every 24 h for a total of 7 days. At the end of the 7-day trial period, the remaining mosquitoes were sexed and counted. Trials were carried out in three biological replicates.

### Statistical Analysis

GraphPad Prism 9 was used for all analyses. Kaplan–Meier curves were plotted to assess differences in population curves. Significant differences were determined using a log-rank Mantel–Cox test with a post hoc analysis and a Bonferroni correction. Parametric, unpaired t-tests were used to identify differences in mean survivorship according to treatment and species (where *α* = 0.01) and to assess the differences between male and female survivorship (where *α* = 0.05)

## Results

### Diols reduce the survivorship of *Ae. aegypti* and *Ae. albopictus* mosquitoes

Of the six diols tested, five led to reduced survivorship in *Ae. aegypti* when compared to mosquitoes that fed on sugar alone (Figure 1). The compounds 1,2-propanediol, 1,3-propanediol, 1,5-pentanediol, 1,6-hexanediol, and 2-methyl-1,3-propanediol yielded reduced mean survivorship over the span of the 7-day trial at a concentration of 5% in a solution of 5% sucrose when compared to a 5% sucrose control, whereas dithiothreitol did not (Table 1). For this portion of the study, we set the alpha value to 0.01 as the P value between the control groups of *Ae. aegypti* and *Ae. albopictus* was 0.039. Of the tested compounds, both 1,2-propanediol and 1,5-pentanediol completely eliminated all adults by the end of the trials, while the other three compounds reduced the survivorship of the population to under 10 %. Additional statistical information is available in Supp. Table 2.

**Fig. 1.**
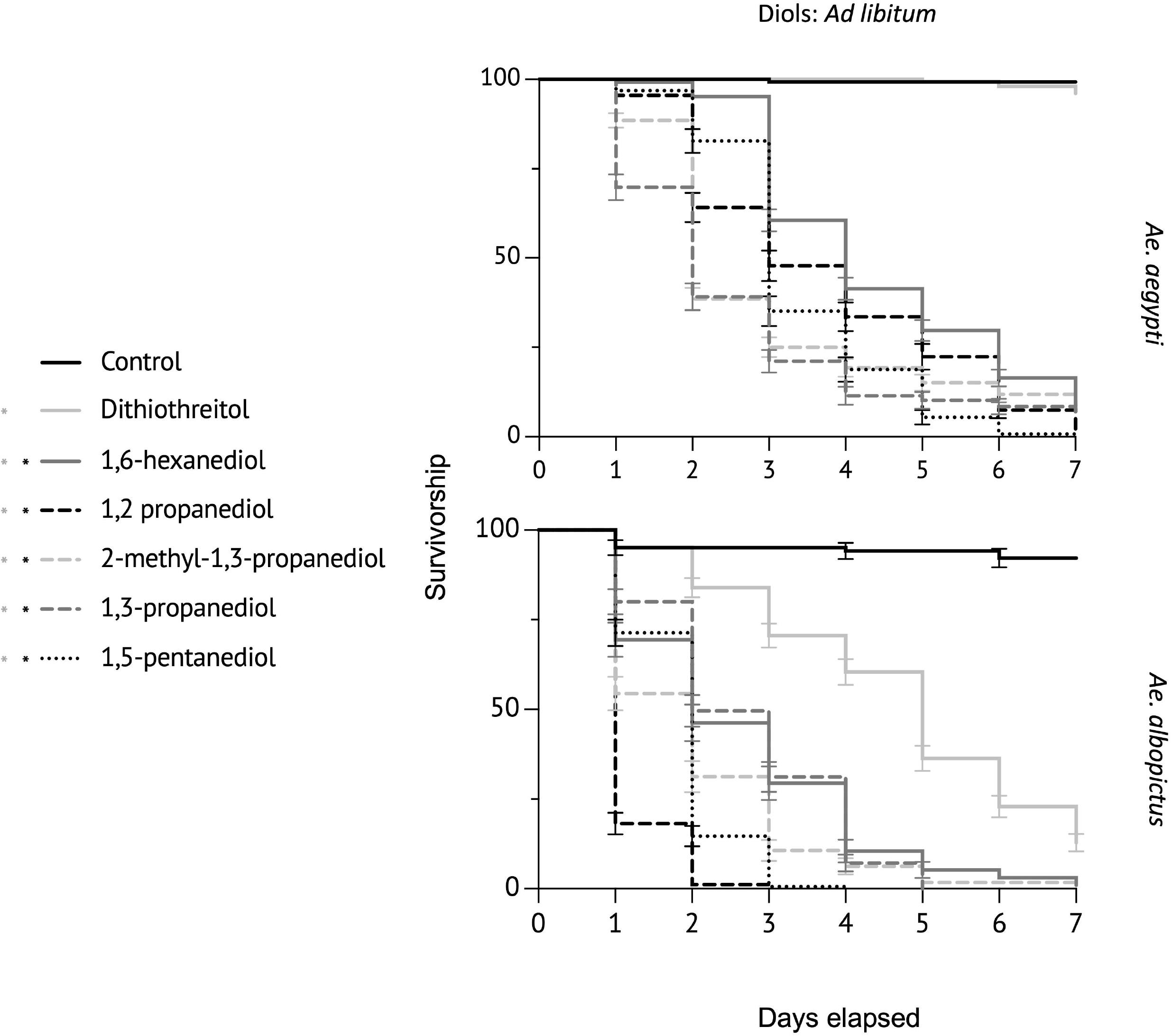

**Table 1.**

| *Ad libitum* trials<br>Compound | *Aedes aegypti*<br>% survive | *Aedes albopictus*<br>% survive | *Aedes aegypti*<br>to control<br>P value | *Aedes albopictus*<br>to control<br>P value | Between<br>Species<br>P value |
| --- | --- | --- | --- | --- | --- |
| Control | 99 | 93 | — | — | 0.0388 |
| 1,2-Propanediol | 0 | 0 | <0.0001 | <0.0001 | — |
| 1,3-Propanediol | 6 | 0 | <0.0001 | <0.0001 | 0.3203 |
| 1,5-Pentanediol | 0 | 0 | <0.0001 | <0.0001 | — |
| 1,6-Hexanediol | 7 | 1 | <0.0001 | <0.0001 | 0.1953 |
| 2-Methyl-1,3-Propanediol | 6 | 1 | <0.0001 | <0.0001 | 0.3712 |
| DL-Dithiothreitol | 97 | 13 | 0.2574 | 0.0007 | 0.0005 |
| Acesulfame Potassium<br>(Ace-K) | 58 | 8 | <0.0001 | 0.0005 | 0.0035 |
| Allulose | 51 | 5 | <0.0001 | <0.0001 | <0.0001 |
| Erythritol | 13 | 6 | <0.0001 | <0.0001 | 0.2534 |
| Monk Fruit Extract | 87 | 79 | 0.0979 | 0.0880 | 0.3858 |
| Neotame | 52 | 6 | 0.0383 | <0.0001 | 0.0480 |
| Sodium Saccharin | 0 | 0 | <0.0001 | <0.0001 | — |
| Stevia | 59 | 15 | 0.0076 | <0.0001 | 0.0055 |
| Sucralose | 45 | 21 | 0.0008 | <0.0001 | 0.0391 |
| Xylitol | 99 | 74 | 0.6874 | 0.1095 | 0.0491 |
| 24-Hour Trials<br>Compound | *Aedes aegypti*<br>% survive | *Aedes albopictus*<br>% survive | *Aedes aegypti*<br>to control<br>P value | *Aedes albopictus*<br>to control<br>P value | Between<br>Species<br>P value |
| Control | 98 | 93 | — | — | 0.0388 |
| 1,2-Propanediol | 42 | 18 | 0.0463 | 0.0014 | 0.3526 |
| 1,3-Propanediol | 63 | 69 | 0.0101 | 0.0022 | 0.4897 |
| 1,5-Pentanediol | 83 | 64 | 0.0310 | 0.0088 | 0.0689 |
| 1,6-Hexanediol | 99 | 44 | 0.7907 | 0.0535 | 0.0367 |
| 2-Methyl-1,3-Propanediol | 78 | 30 | 0.1326 | 0.0005 | 0.0181 |
| Erythritol | 96 | 59 | 0.4535 | 0.2209 | 0.1904 |
| Sodium Saccharin | 90 | 51 | 0.1805 | 0.0189 | 0.0310 |

We then performed the toxicity assays on *Ae. albopictus*, and found that all compounds significantly reduced survivorship compared to sucrose alone (Figure 1). Interestingly, all the diols except for 1,3-propanediol decreased the survivorship curves of *Ae. albopictus* more dramatically (P<0.0001, Log-rank Mantel-Cox), although not the mean survivorship. In *Ae. albopictus*, three compounds, 1,3-propanediol, 1,2-propanediol, and 1,5-pentanediol killed all adults within 7 days. The most significant difference between the two species was the toxicity of DL-dithiothreitol, which did not produce any discernable negative effects on *Ae. aegypti*, but reduced the population of *Ae. albopictus* to 13% (Figure 1).

### Sugar substitutes reduce the survivorship of *Ae. aegypti* and *Ae. albopictus* mosquitoes

Of the sugar substitutes tested on *Ae. aegypti*, all but xylitol yielded changes to survivorship curves (P<0.0001, Log-rank Mantel-Cox) (Figure 2). Compared to the control, however, only acesulfame potassium, allulose, erythritol, sodium saccharin, stevia, and sucralose resulted in a significant reduction of mean survivorship through the course of the seven-day feeding trial (Table 1). The two most toxic substances, erythritol and sodium saccharin, also yielded significantly lower survivorship than the other sugar substitutes, with sodium saccharin as the most potent (Figure 2).

**Fig. 2.**
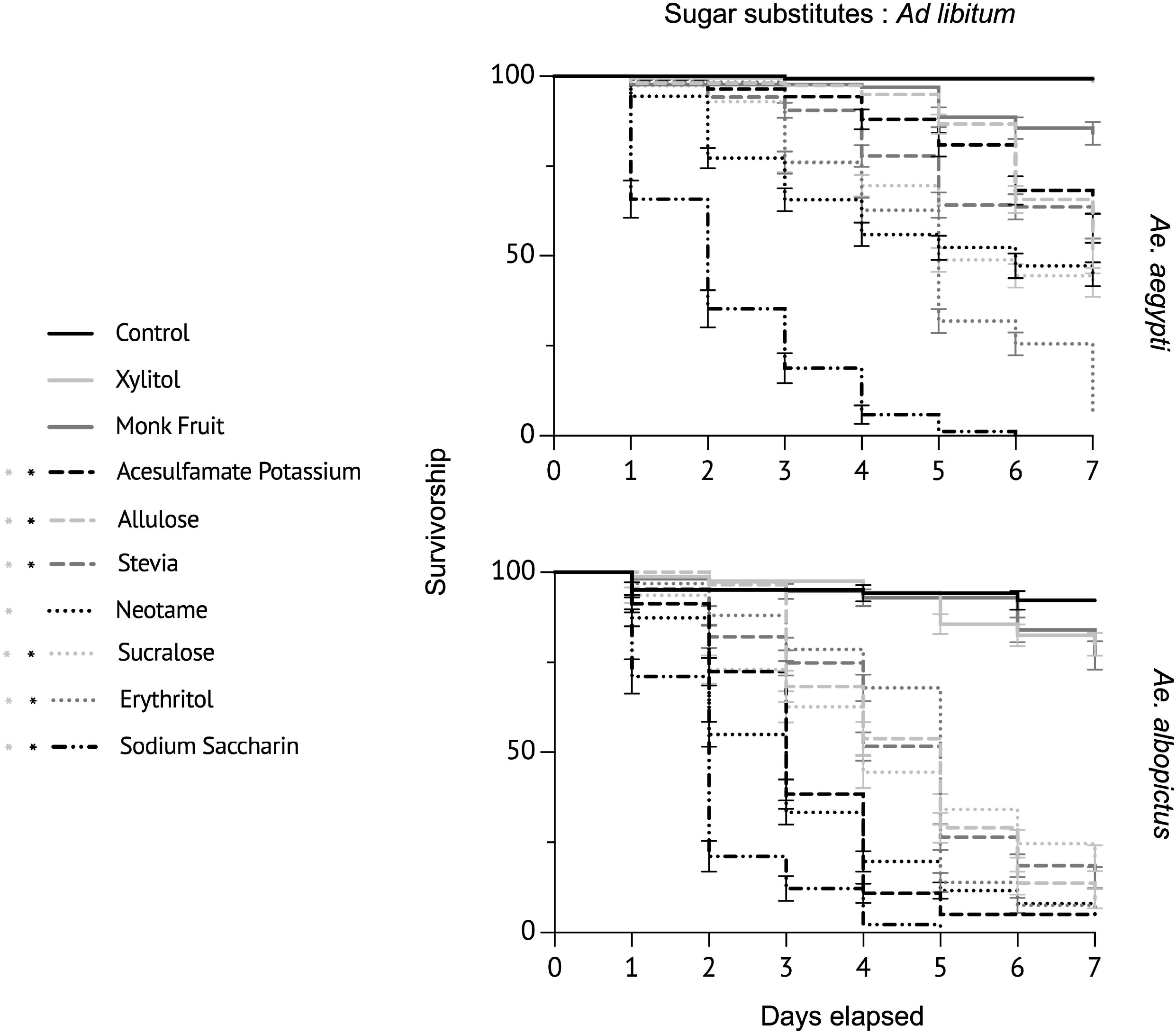

Interestingly, some sugar substitutes exerted more significant effects in *Ae. albopictus* than in *Ae. aegypti*. Acesulfame potassium, allulose, and Stevia yielded significantly lower mean survivorship (Table 1). Lastly, the two compounds that were most potent in *Ae. aegypti* were also among the most toxic compounds in *Ae. albopictus* (Figure 2).

### Toxic compounds reduce the survivorship of *Aedes* mosquitoes after short-term exposure

We then tested the compounds that demonstrated the most toxicity to both species of mosquitoes for their effects after 24 hours of exposure. For *Ae. aegypti*, only 1,3-propanediol reduced the mean survivorship compared to the control(Figure 3), while for *Ae. albopictus*, 1,2-propanediol, 1,3-propanediol, 1,5-pentanediol, and 2-methyl-1,3-propanediol (Table 1, Figure 3). However, for several compounds, the survival curves were significantly different, suggesting that although the mean survivorship did not differ significantly, the rates of population reduction did. Four compounds, 1,2-propanediol, 1,3-propanediol, 2-methyl-1,3-propanediol, and 1,5-pentanediol, changed the survivorship curves of *Ae. aegypti* mosquitoes significantly (P<0.0001, Log-rank Mantel-Cox). Additionally, while the mean survivorship of the two species did not differ, the survivorship curves of *Ae. albopictus* were lower than those of *Ae. aegypti* for five of the compounds tested (1,5-pentanediol, P=0.0005, 1,6-hexanediol, P<0.0001, 2-methyl-1,3-propanediol, P<0.0001, erythritol, P<0.0001, sodium saccharin, P<0.0001 Log rank Mantel-Cox).

**Fig. 3.**
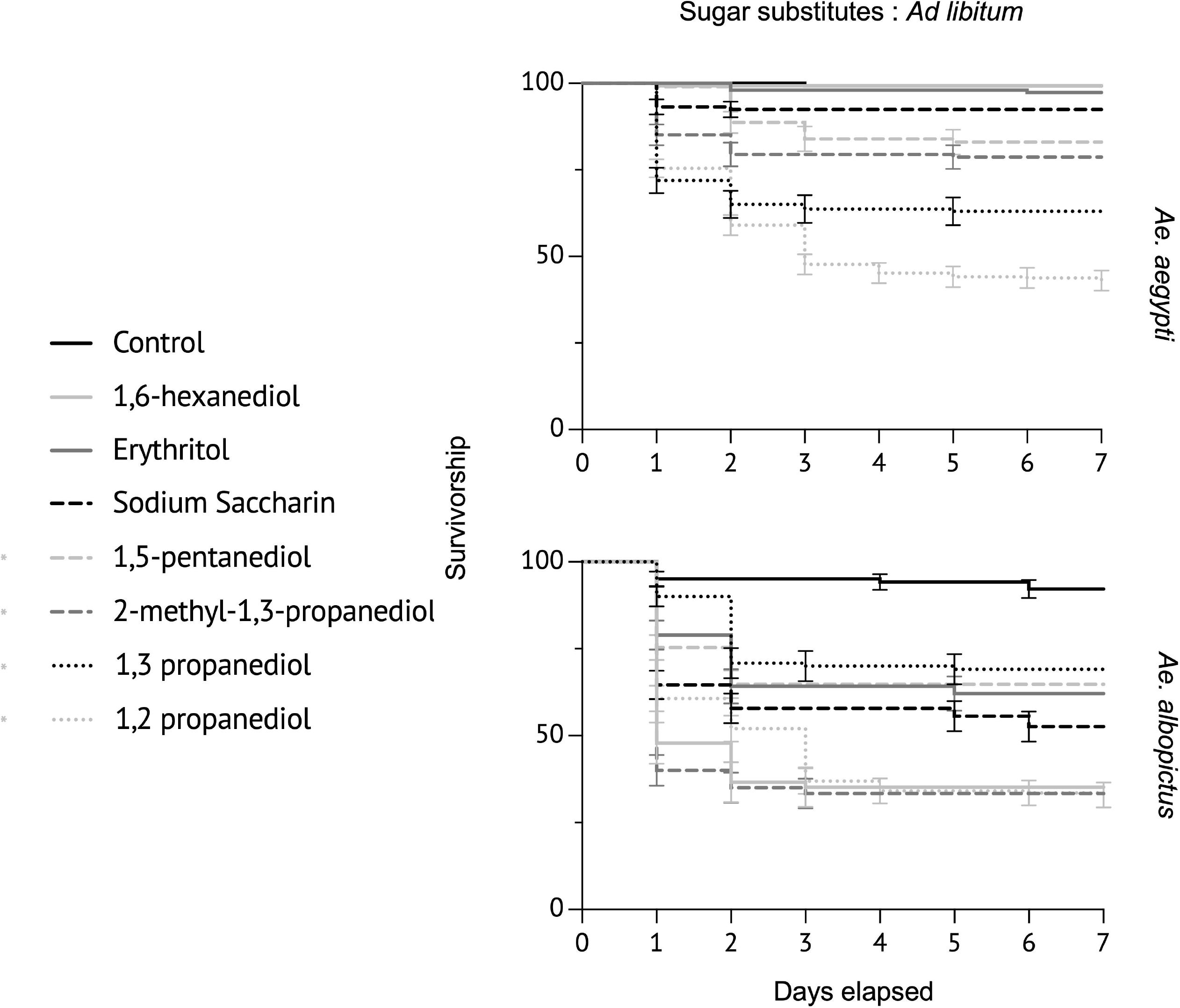

### Toxic compounds have a greater effect on male survivorship than female survivorship

Across species and feeding assay types, the survivorship of male mosquitoes decreased more than that of females. In the *ad libitum* trial, females survived until the end of the experiment in eight conditions, while males survived in only two, and to a lesser degree than the females. In four conditions, females retained significantly higher survivorship means than males: in the *Ae. aegypti ad libitum* trials with erythritol (P=0.0254, unpaired t-test), and in the *Ae. albopictus* 24-hour trials with1,5-pentanediol (P=0.0159, unpaired t-test), 2-methyl-1,3-propanediol (P=0.0030, unpaired t-test), and sodium saccharin (P=0.0307, unpaired t-test) (Figure 4).

**Fig. 4.**
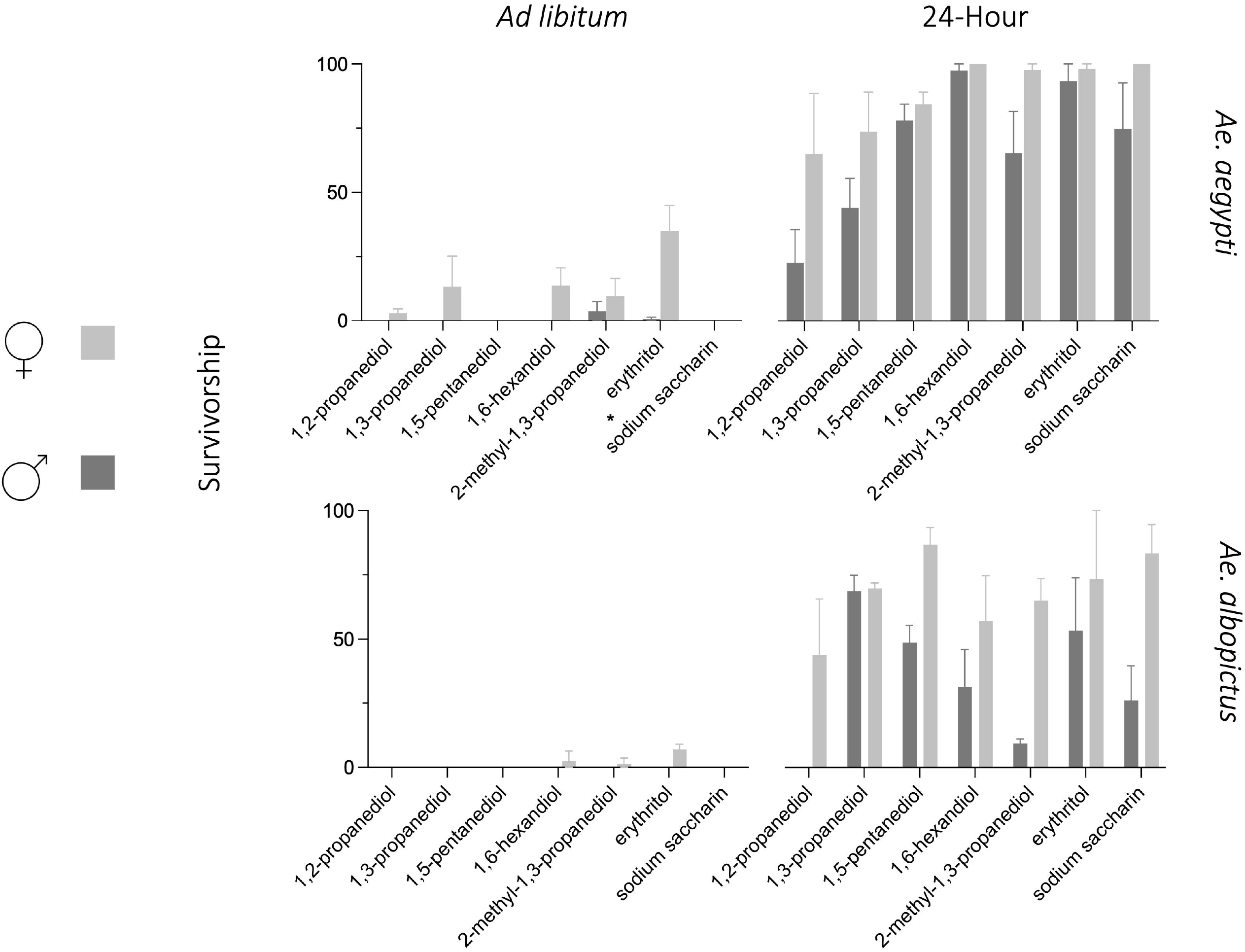

## Discussion

Our results indicate that a variety of diols and sugar substitutes, previously untested for their efficacy as mosquito toxins, reduce the daily survivorship of two major disease vectors: *Ae. aegypti* and *Ae. albopictus*. Decreasing the daily survivorship of adult mosquitoes can yield a decline in population life expectancy and presumably interrupt disease transmission cycles. This outcome could be achieved even at lower, non-fatal concentrations. Decreasing the typical life span of female mosquitoes, even to a moderate degree, could reduce the number of females that survive long enough to complete the extrinsic incubation period for viruses after infection, preventing them from spreading disease from host to host. The non-fatal concentrations may also reduce the likelihood of vectors developing resistance mechanisms as they would likely have reproduced before experiencing the effects of the toxins.

We initially tested the diols and the sugar substitutes on *Ae. aegypti*. Our results indicated that five of the six diols, one of which we previously demonstrated to reduce survivorship, were acutely toxic to mosquitoes and yielded a significant reduction in survivorship by the end of the seven-day trial. While 1,2-propanediol, 1,3-propanediol, 1,5-pentanediol, 1,6-hexanediol, and 2-methyl-1,3-propanediol reduced the population to less than 10% of the starting population, dithiothreitol did not. At the end of the seven-day experiment, we observed no difference in the survivorship of the population that fed on sucrose alone and the population that fed on DL-dithiothreitol mixed with sucrose. The reason for this is yet unexplored, as based on structural similarity and typical commercial use, we hypothesized that all these compounds would prove toxic. Following the conclusion of these trials, we tested the compounds on *Ae. albopictus*. We observed that the compounds that were toxic to *Ae. aegypti* were also harmful to *Ae. albopictus* and dramatically reduced the population within seven days.

Interestingly, dithiothreitol also reduced survivorship. The mean survivorship was lower for all compounds in *Ae. albopictus* than in *Ae. aegypti*, the only compound that significantly reduced that mean survivorship in one species over another was dithiothreitol. This indicates that though these species are closely related evolutionarily and possess overlapping life-histories, key physiological or behavioral differences could be further explored.

We then tested a variety of sugar substitutes, beginning with *Ae. aegypti*. While all substances tested are classified as GRAS, six of the nine compounds produced significant effects on mosquito mean survivorship by the end of the experimental period. Notably, two sugar substitutes - erythritol and sodium saccharin - dramatically reduced the mean survivorship to 13% and 0%, respectively. We then tested the sugar substitutes on *Ae. albopictus*. As with the diols, the sugar substitutes decreased mean survivorship in *Ae. albopictus*. Acesulfame potassium, allulose, and Stevia all yielded significantly lower mean survivorship when compared to *Ae. aegypti*, further indicating that these species may have exploitable physiological differences.

We completed a series of short-term feeding assays for most toxic compounds, in which mosquitoes had access to the toxin mixed with a sucrose solution for 24 hours, at which point it was replaced with a sucrose-only solution. From these trials, we found that four of the seven compounds (1,2-propanediol, 1,3-propanediol, 1,5-pentanediol, and 2-methyl-1,3-propanediol) selected yielded changes to the survivorship curves (P<0.0001, log-rank Mantel-Cox) though did not produce a significant difference in mean survivorship. Though changes in survivorship curves were also evident in *Ae. albopictus*, four of the tested compounds also yielded a significant reduction in mean survivorship. For the short-term trials, 1,2-propanediol produced the most drastic survivorship reduction for both species. Inversely, 1,6-hexanediol significantly reduced the survivorship of *Ae. albopictus*, but not of *Ae. aegypti*. As we initially captured both species in the same geographical region, raised them in the same laboratory conditions, and completed the same experimental treatments, it is plausible that this represents an actual difference in the toxicity of 1,6-hexanediol in one species over another. While *Ae. aegypti* and *Ae. albopictus* have many similarities, including geographic distribution, oviposition choice, and attraction to chemical cues (Kraemer *et al*. 2015, Gunithilaka *et al*. 2017, Hutcheson *et al*. 2022), there may be important differences that allow for the targeting of one species rather than another that require further exploration.

Lastly, we determined that the toxic compounds had more potent effects on the males of both species and in both conditions. This result is likely due to behavioral differences between males and females; male mosquitoes expend more energy in flight and consume sugar more frequently than females, who tend to rest for extended periods.

An essential factor that requires investigation is the potential off-target effects of these compounds. From the significant differences we observed between two closely related mosquito species, we hypothesize that these compounds will have a different impact across all arthropods. This may allow for a more selected and targeted toxin. However, utilizing these compounds in an ATSB system along with attractants and mechanical selection methods could circumvent potential off-target effects. While we initially included aspartame, another widely used sugar substitute, in our study study, we found that its low solubility in water makes it a less practical and accessible ATSB component. We initially tested this compound at a concentration of 1%, as it did not dissolve readily at higher concentrations and saw no effect on mosquito survivorship. However, the toxicity of aspartame to *Ae. aegypti* and *Ae. albopictus* remains an important line of questioning.

Our results indicate that a variety of sugar substitutes and diols reduce the daily survivorship of adult *Ae. aegypti* and *Ae. albopictus* mosquitoes, and by extension, may reduce the average life expectancy of a vector population, leading to a reduction of disease incidence. Many other GRAS compounds, such as potassium sorbate and sodium benzoate (Neeboo 2014), have been used to diminish microbial growth and may be effective against arthropods. With the rising concern of insecticide resistance in mosquitoes worldwide, new and innovative control strategies require prioritization. This study offers insight into the unique compounds that may be used for localized population and disease transmission reduction.

Future field studies integrating an attractive component to the toxic sugar solution will help determine how well these compounds work as ATSB components. Our initial study demonstrates that some of these substances significantly decrease survivorship in two vector mosquito species in a lab setting. We conclude that they may decrease the average life expectancy in natural populations. In the present condition, our study contributes to the groundwork of ATSB design and field-based population studies to locally reduce vector mosquito populations and avoid damage to non-target species.

## Author Contributions

RJP: Conceptualization; Funding acquisition; Resources; Supervision; Project administration; Writing-review and editing. HP: Data curation; Formal analysis; Investigation; Methodology; Supervision; Validation; Visualization; Writing-original draft. ABM: Investigation.

## Acknowledgments

We would like to acknowledge Ben Turnley for assistance in data collection.

This project was funded by the American Mosquito Research Association: AMCARF project number 2021-01.

