## Supplementary material for "Diols and sugar substitutes in attractive toxic sugar baits targeting *Aedes aegypti* and *Aedes albopictus* (Diptera: Culicidae) mosquitoes": Supp. Table 1

| **Diols**  Compound | CAS | Company |
| --- | --- | --- |
| 1,2-Propanediol | 57-55-6 | Fisher Science Education, Nazareth, PA, USA |
| 1,3-Propanediol | 504-63-2 | TCI, Portland, OR, USA |
| 1,5-Pentanediol | 111-29-5 | Acros Organics, Fair Lawn, NJ, USA |
| 1,6-Hexanediol | 629-11-8 | Alfa Aesar, Heysham, Lancashire, UK |
| 2-Methyl-1,3-Propanediol | 2163-42-0 | TCI, Portland, OR, USA |
| DL-Dithiothreitol | 3483-12-3 | bioWORLD, Dublin, OH, USA |
| **Sugar Substitutes**  Compound | CAS | Company |
| Acesulfame Potassium (Ace-K) | __ | Prescribed for Life, Fredericksburg, TX, USA |
| Allulose | __ | 138 Foods, Inc. Claremontm, CA, USA |
| Erythritol | __ | Wholesome Sweeteners, Sugarland, TX, USA |
| Monk Fruit Extract | __ | MarkNature, Fullerton, CA, USA |
| Neotame | __ | MarkNature, Fullerton, CA, USA |
| Sodium Saccharin | 81-07-2 | Loud Wolf, Dublin, CA, USA |
| Stevia | __ | Micro Ingredients, Diamond Bar, CA, USA |
| Sucralose | __ | Bulk Supplements, Henderson, NV, USA |
| Xylitol | __ | Now Foods, Bloomingdale, IL, USA |

Supp. Table 2: Mean survivorship, P values, t values, and degrees of freedom from unpaired t-tests.

*Ad libitum*

trials

Compound

*Aedes aegypti*

% survive

*Aedes albopictus*

% survive

*Aedes aegypti*

to

control P value

*Aedes aegypti*

to

control t,

df

*Aedes albopictus*

to

control P

value

*Aedes albopictus*

to control t,

df

Between

Species

P value

Between Species

t,

df

Control

99

93

__

__

__

__

0.0388

2.44, 2

1,2

-

Propanediol

0

0

**<0.0001**

149.00, 4

**<0.0001**

43.09, 4

N/A

N/A

1,3

-

Propanediol

6

0

**<0.0001**

16.15, 4

**<0.0001**

43.09, 4

0.3203

1.13, 4

1,5

-

Pentanediol

0

0

**<0.0001**

149.00, 4

**<0.0001**

43.09, 4

__

__

1,6

-

Hexanediol

7

1

**<0.0001**

25.49, 4

**<0.0001**

37.81, 4

0.1953

1.55, 4

2

-

Methyl

-

1,3

-

Propanediol

6

1

**<0.0001**

19.84, 4

**<0.0001**

38.98, 4

0.3712

1.01, 4

DL

-

Dithiothreitol

97

13

0.2574

1.32, 4

**0.0007**

9.52, 4

**0.0005**

10.28, 4

Acesulfame Potassium (Ace

-

K)

58

8

**<0.0001**

34.00, 4

**0.0005**

10.14, 4

**0.0035**

6.17, 4

Allulose

51

5

**<0.0001**

36.13, 4

**<0.0001**

30.41, 4

**<0.0001**

20.26, 4

Erythritol

13

6

**<0.0001**

19.55, 4

**<**

**0.0001**

31.00, 4

0.2534

1.33, 4

Monk Fruit Extract

87

79

0.0979

2.15, 4

0.0880

2.25, 4

0.3858

0.97, 4

Neotame

52

6

0.0383

3.04, 4

**<0.0001**

17.48, 4

0.0480

2.82, 4

Sodium Saccharin

0

0

**<0.0001**

149.00, 4

**<0.0001**

43.09, 4

__

__

Stevia

59

15

**0.0076**

4.98, 4

**<0.0001**

29.73, 4

**0.0055**

5.46, 4

Sucralose

45

21

**0.0008**

19.11, 4

**<0.0001**

15.76, 4

0.0391

3.23, 4

Xylitol

99

74

0.6874

0.43, 4

0.1095

2.05, 4

0.0491

2.795, 4

24

-

Hour Trials

Compound

*Aedes aegypti*

% survive

*Aedes albopictus*

% survive

*Aedes aegypti*

to

control P value

*Aedes aegypti*

to

control t,

df

*Aedes albopictus*

to control P

value

*Aedes albopictus*

to control t,

df

Between Species

P value

Between Species

t,

df

Control

98

93

__

__

__

__

0.0388

2.44, 2

1,2

-

Propanediol

42

18

0.0463

2.85, 4

**0.0014**

7.85, 4

0.3526

1.05, 4

1,3

-

Propanediol

63

69

0.0101

4.59, 4

**0.0022**

7.02, 4

0.4897

0.76, 4

1,5

-

Pentanediol

83

64

0.0310

3.26, 4

**0.0088**

4.769, 4

0.0689

2.47, 4

1,6

-

Hexanediol

99

44

0.7907

0.28, 4

0.0535

2.71, 4

0.0367

3.09, 4

2

-

Methyl

-

1,3

-

Propanediol

78

30

0.1326

1.89, 4

**0.0005**

10.11, 4

0.0181

3.86, 4

Erythritol

96

59

0.4535

0.83, 4

0.2209

1.45, 4

0.1904

1.57, 4

Sodium Saccharin

90

51

0.1805

1.62, 4

0.0189

3.82, 4

0.0310

3.26, 4
